# Computational Identification of Candidate Gene Families for Volatile Sulfur Compound Biosynthesis in *Cannabis sativa* Using Profile Hidden Markov Models

**DOI:** 10.64898/2026.08.18.745489

**Authors:** Emanuel Maminakis, Logan Geffen, Kevelin Barbosa-Xavier, Suliman Sharif

## Abstract

Cannabis is well known for its pungent, skunk-like aroma. Recent chemical studies have identified prenylated and C6 volatile sulfur compounds as contributors to its skunky and citrus-like aromas, but the pathways that produce these compounds remain unknown. This gap limits efforts to explain variation in sulfur-aroma traits and to selectively enhance or reduce those traits. To address this gap, we used the known chemistry of sulfur-containing volatiles in Cannabis and characterized sulfur and volatile biosynthetic pathways in other plant species to select candidate enzyme groups. Because the GMO cultivar is anecdotally associated with a pronounced sulfurous aroma, reference protein sequences and profile hidden Markov models were used to search its version 1 (v1) primary high-confidence protein set of 55,790 sequences. These searches recovered 975 unique proteins. Sequence screening retained 941 candidates across 20 reporting categories; 939 contained all expected domains, while the two candidates assigned to the methionine gamma-lyase (MGL)-nearest category had no category-specific expected-domain rule. The largest reporting category comprised 359 proteins containing a cytochrome P450 domain, recovered through a search motivated by cytochrome P450 family 74 (CYP74) enzymes involved in oxylipin and plant volatile formation. Thirteen of these proteins were also recovered by at least one full-length CYP74 reference search. Other large reporting categories included 218 sugar-transferase, 83 glutathione-transferase, and 61 alcohol dehydrogenase candidates. Comparison with the Cannabis Expression Atlas linked 168 candidates to 128 annotated genes through 100%-identity amino-acid matches spanning at least 80% of each GMO v1 candidate protein. Twenty-nine genes were tissue-specific, including 13 root-specific and 6 trichome-specific genes. These results define candidates for biochemical testing and direct searches for additional enzymes acting upstream and downstream in Cannabis sulfur-volatile pathways.

## 1. Introduction

The aroma of Cannabis is a major part of how cultivars and products are perceived. Among its most recognizable features is the pungent, skunk-like character of many cultivars. Because aroma preferences and product requirements differ, breeders and producers may seek to preserve or intensify this trait where it contributes to consumer appeal, or reduce it in plants and products intended to have less odor. Isolating or producing the responsible compounds could also support analytical standards and sensory studies. To better understand these traits, recent analytical studies have begun to identify the compounds responsible. Comprehensive two-dimensional gas chromatography identified a family of prenylated volatile sulfur compounds (VSCs), designated VSC3 through VSC7, and showed that 3-methyl-2-butene-1-thiol is a major contributor to the characteristic skunk-like aroma of selected samples (Oswald et al., 2021). A later survey identified a second, C6 class that included 3-mercaptohexanol and its acetate and butyrate esters and linked these compounds to citrus-like aroma differences among cultivars (Oswald et al., 2023). Together, these studies establish at least two structurally distinct classes of aroma-active Cannabis sulfur compounds.

The biosynthetic origins of these compounds remain unresolved. Identifying their precursors and enzymes is needed to explain how sulfur-aroma traits arise and vary in Cannabis, define targets for biochemical validation, and ultimately provide a basis for targeted manipulation of the underlying pathways. No protein family or biosynthetic pathway has yet been directly characterized for VSC production in Cannabis. We therefore turned to better-characterized sulfur-volatile pathways in other plants as models for possible Cannabis biosynthetic routes.

In *Vitis vinifera* (grapevine), glutathione and cysteine conjugates serve as nonvolatile precursors of C6 varietal thiols (Kobayashi et al., 2011). Glutathione S-transferase (GST) activity can form glutathionyl precursors from C6 aldehydes, followed by processing toward cysteinyl conjugates through gamma-glutamyl transferase (GGT) activity and related steps (Kobayashi et al., 2011; Philips et al., 2019). During wine fermentation, the yeast beta-lyase Irc7 cleaves cysteine conjugates to release volatile thiols (Cordente et al., 2019). This route provides a working model for Cannabis C6 sulfur volatiles.

Other organisms provide additional sulfur-release pathways. In *Allium*, alliinase cleaves S-alk(en)yl cysteine sulfoxides, and onion lachrymatory-factor synthase acts on a resulting sulfenic-acid intermediate (Weiner et al., 2009; Masamura et al., 2012). In Brassicaceae, myrosinases hydrolyze glucosinolate precursors to initiate characteristic sulfur chemistry (Oloyede et al., 2021). These pathways use substrates not established in Cannabis, so they were used to identify potentially relevant enzyme families rather than treated as direct models of Cannabis sulfur metabolism. Plant oxylipin metabolism provides a separate model for generating the C6 carbon backbone that could feed a conjugate-based route (Wan et al., 2013). Together, these systems motivate a broad but explicitly hypothesis-based candidate search.

Guided by chemical studies of sulfur-containing volatiles in Cannabis, we examined better-characterized sulfur and volatile biosynthetic pathways in other plant species and selected enzyme families representing plausible biosynthetic steps. The GMO cultivar is a commercial production clone derived from a ChemD × Girl Scout Cookies cross (Lynch et al., 2025). Its v1 proteome was selected because anecdotal reports characterize the cultivar as unusually sulfurous, making it a relevant starting point for a sulfur-volatile candidate survey. Using reference enzyme sequences and profile hidden Markov models (HMMs) as queries, we searched this proteome with phmmer and hmmsearch to identify candidate homologs with the expected domain architectures. The search panel represented candidate enzyme groups associated with sulfur amino-acid and glutathione metabolism; conjugation, transfer, and redox reactions; sulfur release and related reference pathways; and oxylipin and volatile-product biosynthesis. For closely related enzyme families that could not be distinguished by domain architecture alone, candidates were first classified using phylogenetic analysis. Candidate proteins were subsequently screened for conserved sequence motifs and expected domain composition. The retained candidates were then linked to tissue-resolved expression profiles from the Cannabis Expression Atlas.

## 2. Materials and methods

### 2.1 Proteome and candidate-search panel

Protein models for the GMO cultivar were obtained from the Cannabis pangenome resource (Lynch et al., 2025). Searches used its version 1 (v1) primary high-confidence protein set of 55,790 sequences, and full-length sequences were recovered by exact GMO v1 identifier from the accompanying all-protein FASTA. All candidate counts refer to this GMO v1 proteome.

The candidate-search panel comprised prespecified enzyme activities drawn from the comparative pathways. Full-length queries came from the reviewed UniProtKB/Swiss-Prot entries in Table 1, selected as functionally annotated plant representatives of those activities and biochemical contexts; these sequence queries were distinct from the Pfam profile HMMs, whose accessions and model versions are reported in the deposited query-panel manifest (Mistry et al., 2021). Here, a target enzyme group denotes an activity or enzyme-family hypothesis, whereas its search target comprises the listed Pfam profile or domain architecture and any listed full-length phmmer query. All groups were searched with their Pfam HMMs and, where available, the full-length reference. The groups were organized into four biochemical contexts: sulfur amino-acid and glutathione metabolism; conjugation, transfer, and redox reactions; sulfur release and related reference pathways; and oxylipin and volatile-product biosynthesis (Table 1; Figure 1). Cystathionine beta-lyase (CBL), cystathionine gamma-synthase (CGS), methionine gamma-lyase (MGL), and cysteine-S-conjugate beta-lyase-like (CSCBL-like) all used PF01053; their shared candidate pool was resolved jointly to nearest reference groups by phylogenetic distance. Thus, the 23 target enzyme groups in Table 1 yielded 20 final reporting categories (Section 3.4).

**Table 1.** Candidate-search panel: selection rationales, target enzyme groups, reference organisms and UniProt identifiers, and expected Pfam domain architectures.

| Specific selection rationale or source analogy | Broad biochemical context | Target enzyme group | Reference organism | Full-length phmmer reference (UniProt accession/entry name) | Expected Pfam domain architecture |
| --- | --- | --- | --- | --- | --- |
| Methionine-pool target: S-adenosylmethionine synthase (SAMS), which forms S-adenosylmethionine as an activated methionine metabolite | Sulfur amino-acid and glutathione metabolism | SAMS | *Elaeagnus umbellata* (autumn olive) | Q9AT56/METK1_ELAUM | PF00438 + PF02772 + PF02773 |
| Arabidopsis sulfur-amino-acid comparison: cystathionine cleavage in methionine biosynthesis | Sulfur amino-acid and glutathione metabolism | CBL | *Arabidopsis thaliana* (thale cress) | P53780/METC_ARATH | PF01053 |
| Arabidopsis sulfur-amino-acid comparison: entry of cysteine-derived sulfur into methionine biosynthesis | Sulfur amino-acid and glutathione metabolism | CGS | *Arabidopsis thaliana* (thale cress) | P55217/CGS1_ARATH | PF01053 |
| Grape C6-thiol analogy: glutamate-cysteine ligase (GSH1), the first glutathione-biosynthesis step supplying the GST conjugation route (Kobayashi et al., 2011; Jez et al., 2004) | Sulfur amino-acid and glutathione metabolism | GSH1 | *Medicago truncatula* (barrel medic) | Q9ZNX6/GSH1_MEDTR | PF04107 |
| Grape C6-thiol analogy: glutathione synthetase (GSH2), the second glutathione-biosynthesis step supplying the GST conjugation route (Kobayashi et al., 2011; Ullmann et al., 1996) | Sulfur amino-acid and glutathione metabolism | GSH2 | *Brassica juncea* (brown mustard) | O23732/GSHB_BRAJU | PF03199 + PF03917 |
| Cysteine-pool target: O-acetylserine(thiol)lyase (OASTL)-dependent cysteine synthesis supplying glutathione and cysteine conjugates | Sulfur amino-acid and glutathione metabolism | OASTL | *Arabidopsis thaliana* (thale cress) | P47999/CYSKP_ARATH | PF00291 |
| Grape C6-thiol analogy: grapevine GST-mediated formation of glutathionyl precursors from trans-2-hexenal (Kobayashi et al., 2011) | Conjugation, transfer, and redox reactions | GST | *Vitis vinifera* (grapevine) | — | PF02798 |
| Allium and grape precursor-processing analogy: gamma-glutamyl removal from glutathione conjugates toward cysteine conjugates (Philips et al., 2019) | Conjugation, transfer, and redox reactions | GGT | *Allium sativum* (garlic) | — | PF01019 |
| Brassicaceae comparison: uridine diphosphate (UDP)-dependent glycosyltransferase (UGT) activity in precursor-storage chemistry | Conjugation, transfer, and redox reactions | UGT | *Arabidopsis thaliana* (thale cress) | O48676/U74B1_ARATH | PF00201 |
| Brassicaceae comparison: 3'-phosphoadenosine 5'-phosphosulfate-dependent sulfation by sulfotransferase (SULT) enzymes in sulfur-conjugation chemistry | Conjugation, transfer, and redox reactions | SULT | *Arabidopsis thaliana* (thale cress) | Q9C9D0/SOT16_ARATH | PF00685 |
| Brassicaceae comparison: flavin-containing monooxygenase (FMO)-mediated sulfur oxidation of precursor compounds | Conjugation, transfer, and redox reactions | FMO | *Arabidopsis thaliana* (thale cress) | Q9SXE1/GSOX3_ARATH | PF00743 |
| Arabidopsis sulfurtransferase 18 (STR18) comparison: sulfane-sulfur transfer | Conjugation, transfer, and redox reactions | Rhodanese | *Arabidopsis thaliana* (thale cress) | Q9FKW8/STR18_ARATH | PF00581 |
| Wine-yeast analogy: Irc7-like cleavage of cysteine-S-conjugates to release varietal thiols (Cordente et al., 2019) | Sulfur release and related reference pathways | CSCBL-like | *Saccharomyces cerevisiae* (baker's yeast) | — | PF01053 |
| Jackfruit comparison: methionine-gamma-lyase chemistry as a route from methionine to methanethiol | Sulfur release and related reference pathways | MGL-Artocarpus | *Artocarpus heterophyllus* (jackfruit) | — | PF01053 |
| Within-species comparison: annotated Cannabis MGL-like references for the methionine-cleavage hypothesis | Sulfur release and related reference pathways | MGL-Cannabis | *Cannabis sativa* (cannabis) | — | PF01053 |
| Durian comparison: methionine-gamma-lyase chemistry as a sulfur-volatile release route | Sulfur release and related reference pathways | MGL-Durio | *Durio zibethinus* (durian) | — | PF01053 |
| Garlic analogy: cleavage of S-alk(en)yl cysteine sulfoxides to initiate Allium volatile-sulfur chemistry (Weiner et al., 2009; Masamura et al., 2012) | Sulfur release and related reference pathways | Alliinase | *Allium sativum* (garlic) | — | PF04863 + PF04864 |
| Onion analogy: conversion of an alliinase-derived sulfenic-acid intermediate to lachrymatory factor (Masamura et al., 2012) | Sulfur release and related reference pathways | Lachrymatory-factor synthase | *Allium cepa* (onion) | — | PF10604 |
| Brassicaceae analogy: hydrolysis of glucosinolate-type sulfur-storage compounds (Oloyede et al., 2021) | Sulfur release and related reference pathways | Myrosinase | *Armoracia rusticana* (horseradish) | — | PF00232 |
| C6-thiol carbon-backbone context: formation of fatty-acid hydroperoxides by lipoxygenase (LOX), upstream of hydroperoxide lyase (HPL)-derived C6 aldehydes | Oxylipin and volatile-product biosynthesis | LOX | *Humulus lupulus* (hops) | — | PF00305 + PF01477 |
| Arabidopsis cytochrome P450 family 74 (CYP74) comparison: HPL production of C6 aldehydes and allene oxide synthase (AOS) diversion of fatty-acid hydroperoxides toward jasmonate biosynthesis (Song et al., 1993; Wan et al., 2013) | Oxylipin and volatile-product biosynthesis | AOS/HPL-motivated P450 search | *Arabidopsis thaliana* (thale cress) | Q96242/CP74A + Q9ZSY9/C74B2 | PF00067 |
| AOS-branch comparison: allene oxide cyclase (AOC)-mediated cyclization in jasmonate biosynthesis | Oxylipin and volatile-product biosynthesis | AOC | *Arabidopsis thaliana* (thale cress) | Q9LS01/AOC3_ARATH | PF06351 |
| Plant C6-volatile analogy: alcohol dehydrogenase (ADH)-mediated reduction of HPL-derived aldehydes to the corresponding alcohols | Oxylipin and volatile-product biosynthesis | ADH | *Passiflora edulis* (passion fruit) | — | PF00107 + PF08240 |

**Figure 1.**
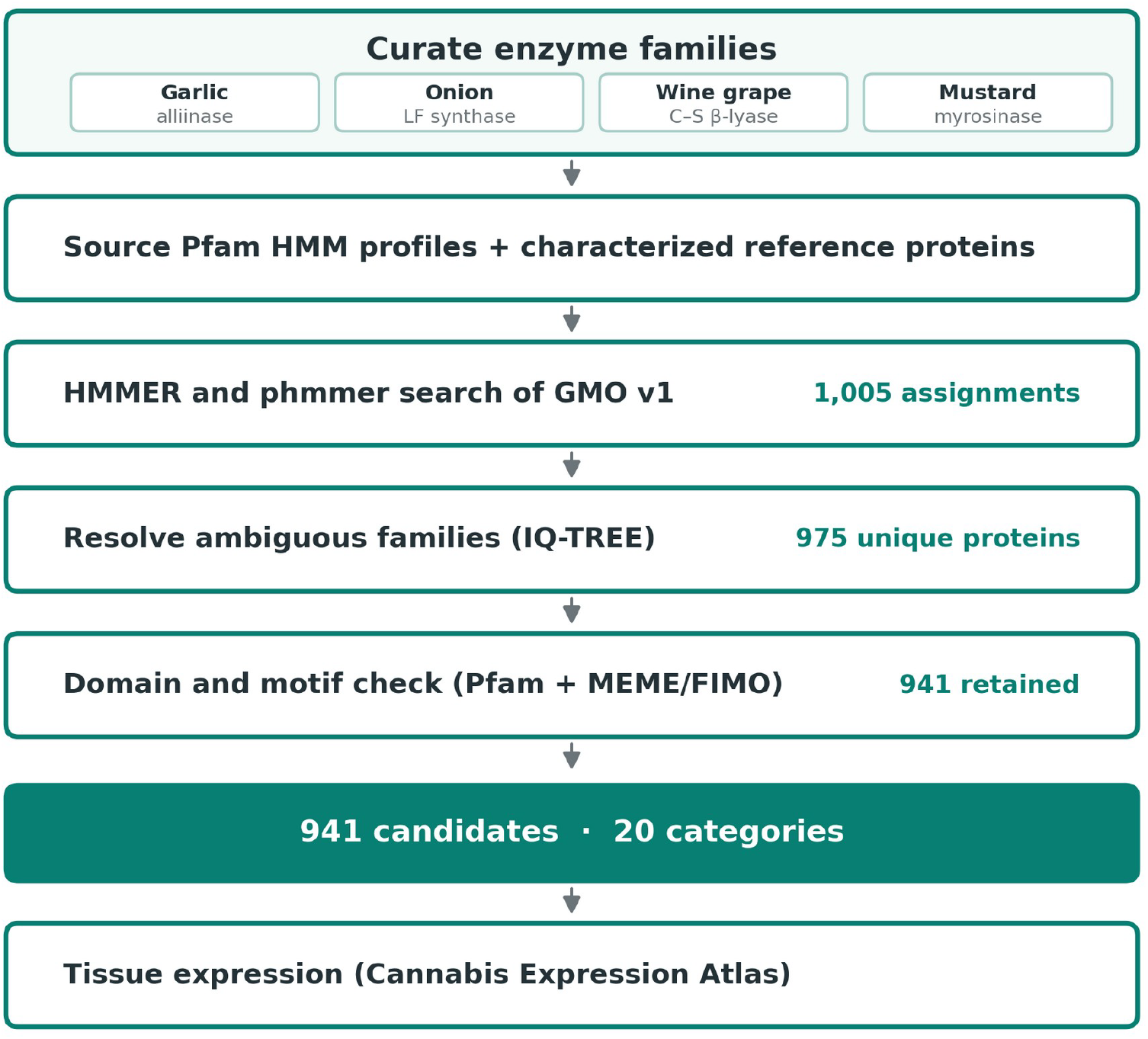
Candidate-search workflow. The candidate-search panel supplied Pfam profiles and, where available, characterized reference proteins for hmmsearch and phmmer searches against the GMO v1 proteome. Shared candidate pools that could not be resolved by domain architecture alone were classified phylogenetically. Candidates were then screened for conserved sequence motifs and expected domain composition before retained proteins were cross-referenced to the Cannabis Expression Atlas.

### 2.2 Pfam-profile searches

Pfam-profile and full-length-reference searches used HMMER 3.4 (Eddy, 2011), with hmmsearch for Pfam profiles and phmmer for full-length protein queries. Every target enzyme group was searched against the 55,790-sequence GMO v1 primary high-confidence proteome with the Pfam HMMs listed in Table 1. Searches used an inclusion threshold of 0.001 and explicitly set -Z to 567,483, which defined the target-sequence count used for per-sequence E-value calculation. Candidate extraction required a full-sequence score of at least 50 and an E-value of at most 1 × 10^−5^. All score-qualified profile hits had reported E-values at or below 5.2 × 10^−12^, so using the input sequence count for E-value calculation would not have changed candidate retention under these thresholds. For target enzyme groups represented by more than one Pfam profile, a protein entered the candidate set if it passed any associated profile.

### 2.3 Full-length-reference searches and candidate-set integration

For target enzyme groups with a full-length reference listed in Table 1, the GMO v1 proteome was searched independently by phmmer. These searches used an inclusion E-value of 0.001 with the default target space, and candidate extraction applied the same score and E-value thresholds as the Pfam searches. A protein entered the downstream workflow if it qualified in either the Pfam-profile or full-length-reference search; recovery by both was recorded as overlap but not required. All resulting candidates were subsequently screened for the expected Pfam domain or, for multidomain groups, the complete listed architecture. The MGL-nearest reporting category was the exception because no category-specific expected-domain rule was defined (Section 2.5). In the AOS/HPL-motivated analysis, a single PF00067 domain-profile search was paired with separate full-length phmmer searches using the Arabidopsis reference proteins Q96242/CP74A and Q9ZSY9/C74B2. Candidates recovered against at least one reference and against both references were counted separately.

### 2.4 Phylogenetic assignment of the PF01053 candidates

The PF01053 profile, used for the CBL, CGS, MGL, and cysteine-S-conjugate-beta-lyase-like target enzyme groups, recovered six unique GMO v1 proteins. Candidate and reference sequences were aligned with MAFFT using automatic strategy selection, trimmed with trimAl in automated mode, and analyzed by maximum likelihood with IQ-TREE 3.1.1 (Katoh et al., 2002; Capella-Gutierrez et al., 2009; Wong et al., 2026). The LG+R2 substitution model was selected by Bayesian information criterion. Branch support was estimated with 1,000 ultrafast bootstrap replicates (Hoang et al., 2018). The reference panel contained 41 UniProt entries spanning the four groups, including seven reviewed proteins. Each GMO candidate was assigned to the reference group with the shortest patristic distance and reported as CBL-nearest, CGS-nearest, or MGL-nearest.

### 2.5 Motif and expected-domain screening

Motif screening was performed separately within each reporting category. For each category, MEME learned up to 10 recurring protein motifs from its GMO v1 candidate sequences, using motif widths of 6 to 50 residues and the zero-or-one occurrence per sequence model (Bailey and Elkan, 1994). FIMO then scanned the same candidate sequences for those MEME-derived motifs at p ≤ 1 × 10^−4^ (Grant et al., 2011). For each sequence, motif coverage was the fraction of that category’s MEME-discovered motifs detected in the sequence; it measured sequence consistency within each candidate group. Candidates passed motif screening if they met either a primary or partial motif criterion. The default primary criterion required at least two distinct motifs at a coverage of at least 0.20; ADH, GST, and UGT required at least three motifs and 0.25 coverage, and the alliinase/1-aminocyclopropane-1-carboxylate (ACC)-synthase-like category required at least two motifs and 0.25 coverage. The default partial criterion required at least one motif and 0.10 coverage, or 0.12 coverage for the alliinase/ACC-synthase-like category. Because GSH2 had only one candidate, that candidate proceeded directly to expected-domain screening without within-category motif discovery.

Each candidate was then rescanned with hmmscan against the Pfam profiles specified for its category (Mistry et al., 2021), applying a full-sequence E-value threshold of 1 × 10^−5^ (-E 1e-5) and a domain E-value threshold of 1 × 10^−3^ (--domE 1e-3). This final gate required the candidate to contain the complete listed architecture of the target Pfam profile. The MGL-nearest reporting category had no category-specific expected-domain rule. Its two members, GMO.00001695.v1.g225230.t1 and GMO.00002556.v1.g618800.t1, were therefore retained without a category-specific hmmscan result. The other 939 retained proteins each have a recorded complete-architecture result. Stability, contamination, physicochemical, paralog, and PROSITE motif-match fields were retained as annotations and were not used to remove candidates.

### 2.6 Cannabis Expression Atlas cross-reference

To connect the 941 retained GMO v1 proteins with Cannabis Expression Atlas genes, we analyzed their BLASTp matches to atlas proteins (Altschul et al., 1997). BLASTp used BLOSUM62, E-value ≤ 0.001, query coverage per high-scoring segment pair (HSP) ≥ 80%, and one target sequence per query. For each GMO protein, the highest-bitscore HSP was retained. A primary mapping required 100% amino-acid identity in that HSP and coverage of at least 80% of the full GMO protein. Atlas gene classification, tissue assignment, and Tau values were then joined to the matched subject identifier (Barbosa-Xavier et al., 2024).

Each GMO candidate-to-atlas hit was counted as one candidate mapping, and tissue summaries collapsed atlas subject identifiers so that a gene matched by multiple GMO candidates was counted only once across tissues. To measure sensitivity to sequence identity, non-primary mappings were also counted from 80% through 100% identity while maintaining at least 80% query coverage.

### 2.7 Use of large language models

Large language models accessed through Anthropic Claude, OpenAI Codex, and Google Gemini were used to assist with drafting and troubleshooting analysis code and developing scripts used to verify processed results and manuscript claims against source data. The authors defined the analyses, supplied the underlying data and scientific logic, reviewed the generated code, and verified all reported results against the analysis outputs.

## 3. Results

### 3.1 Profile and reference searches recovered 975 unique proteins

Combining qualifying Pfam-profile and full-length-reference hits by target enzyme group produced 1,005 search-level candidate assignments representing 975 unique GMO v1 proteins (Figure 2A). The PF01053 profile generated 36 assignments for six unique proteins across the CBL, CGS, cysteine-S-conjugate beta-lyase-like, and three MGL target enzyme groups, so 30 assignments were repeats. All 975 unique proteins underwent sequence screening; the six proteins in the shared PF01053 pool also underwent phylogenetic assignment.

**Figure 2.**
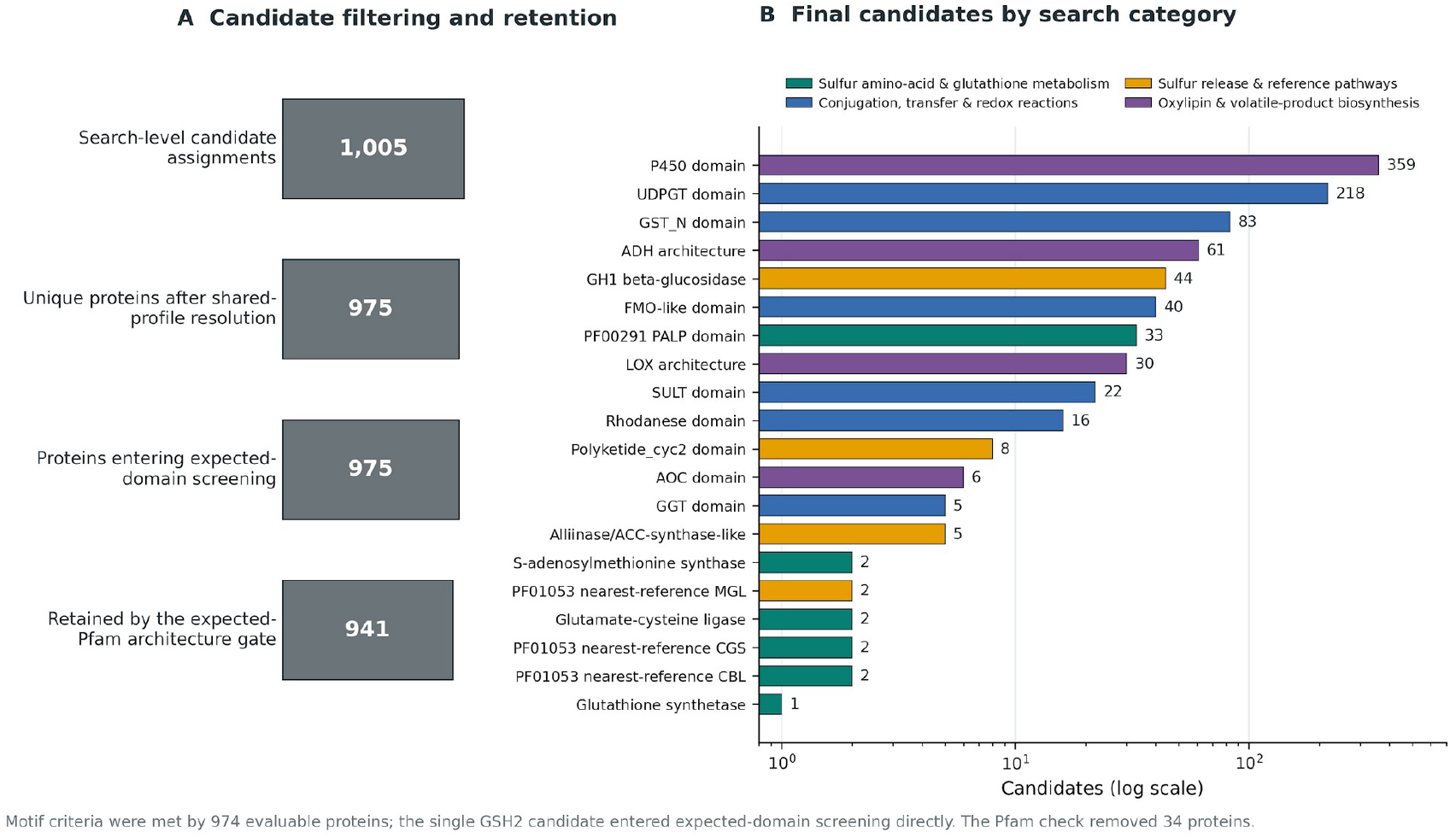
Candidate filtering and final reporting-category counts. (A) Search assignments, unique proteins after shared-profile resolution, and retained proteins. Expected domain architectures were recorded for 939 proteins; two MGL-nearest proteins were retained without a category-specific expected-domain rule. (B) Retained proteins by reporting category on a logarithmic scale. In panel B, colors denote sulfur amino-acid and glutathione metabolism (teal), conjugation, transfer, and redox reactions (blue), sulfur release and related reference pathways (orange), and oxylipin and volatile-product biosynthesis (purple).

### 3.2 Full-length-reference searches contributed 25 retained candidates

Among retained target enzyme groups with full-length references, the full-length-reference searches added 27 proteins not recovered by the corresponding Pfam profile: 24 UGT, one SULT, and two FMO candidates. Every other reference hit was also recovered by its Pfam profile. Expected-domain screening removed the two FMO candidates because they lacked PF00743, leaving 25 retained proteins that entered through the full-length-reference route.

The PF00067 profile recovered 359 proteins, including all 13 proteins recovered by at least one of the two Arabidopsis CYP74 references; 11 were recovered by both. PF00067 models the conserved cytochrome P450 domain, so all 359 proteins were retained as a broad P450-domain candidate set.

### 3.3 Phylogenetic distance assigned six PF01053 candidates to reference groups

Shortest patristic distance in the maximum-likelihood phylogeny placed two PF01053 candidates nearest the CBL reference group, two nearest CGS, and two nearest MGL (Figure 3). No candidate was placed nearest the cysteine-S-conjugate-beta-lyase-like group.

**Figure 3.**
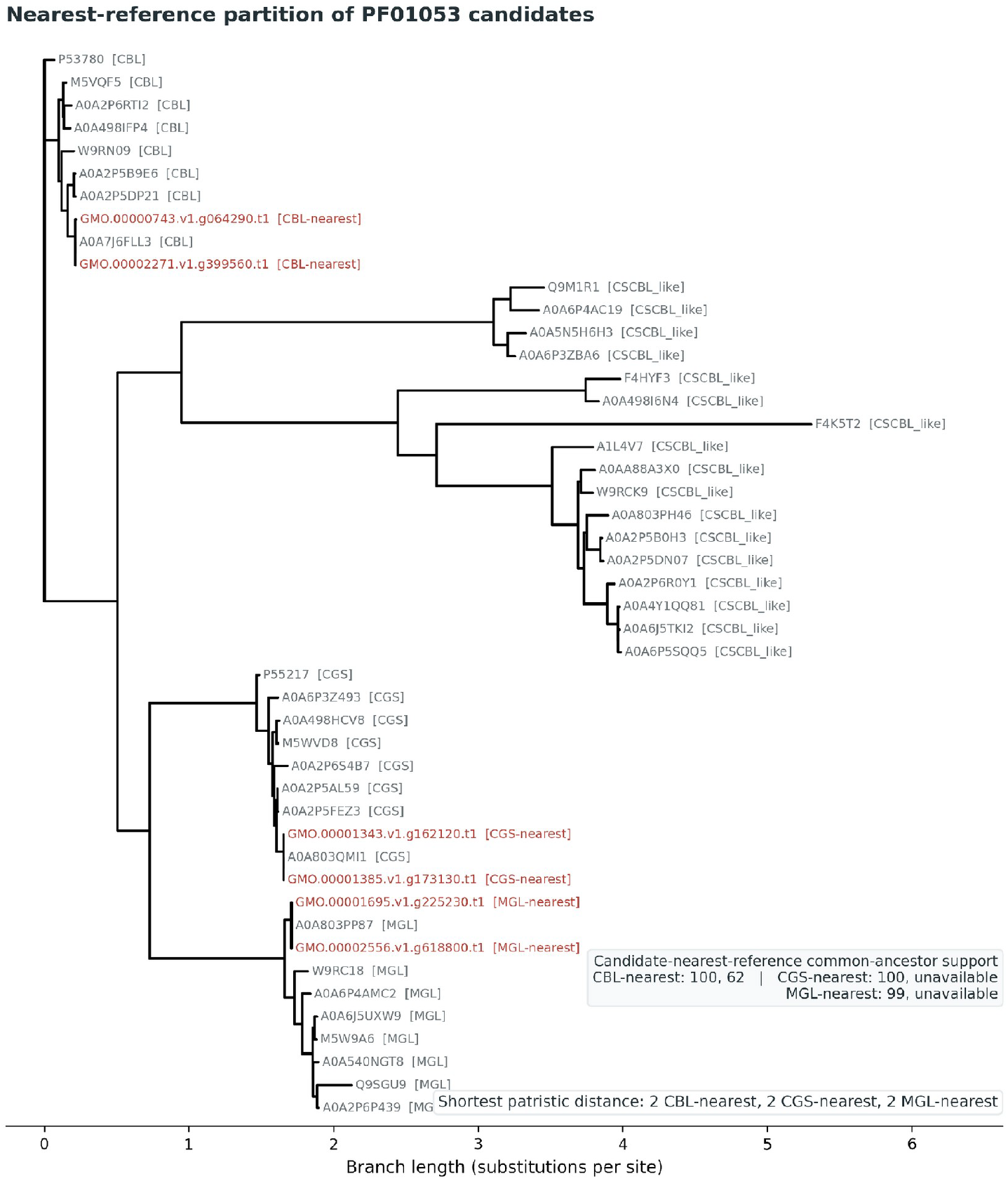
Nearest-reference partition of PF01053 candidates. The maximum-likelihood tree includes six GMO v1 candidates and CBL, CGS, MGL, and cysteine-S-conjugate-beta-lyase-like references. Shortest patristic distance assigned two candidates to each of the CBL-, CGS-, and MGL-nearest groups. Candidates are shown in red; branch lengths indicate substitutions per site. Assignments did not use a support cutoff.

### 3.4 Sequence screening retained 941 candidates across 20 reporting categories

MEME/FIMO evaluated 974 of the 975 proteins, and all 974 met the motif criteria. The single GSH2 candidate proceeded directly to expected-domain screening. The expected-Pfam screen removed 34 proteins: 22 with incomplete ADH architecture, 5 with incomplete alliinase/ACC-synthase-like architecture, 2 lacking the FMO-like domain, and 5 with incomplete LOX architecture. The final catalog contained 941 proteins across 20 reporting categories, corresponding to 96.5% of the proteins entering domain screening (Figure 2A). Of the retained proteins, 939 have a recorded complete expected-domain result; the two MGL-nearest proteins were retained because no category-specific expected-domain rule was defined (Section 2.5).

The four biochemical contexts contained 42 candidates related to sulfur amino-acid and glutathione metabolism, 384 related to conjugation, transfer, and redox reactions, 59 related to sulfur release and related reference pathways, and 456 related to oxylipin and volatile-product biosynthesis. The largest reporting categories were the PF00067 cytochrome P450 domain with 359 candidates, the PF00201 UDP-glycosyltransferase domain with 218, the PF02798 GST_N domain with 83, and the PF00107+PF08240 ADH architecture with 61 (Figure 2B).

Table 2 gives the count for every reporting category and its relationship to the pre-search design. Seventeen of the 23 target enzyme groups in Table 1 map to one reporting category each. The remaining six target enzyme groups—CBL, CGS, cysteine-S-conjugate beta-lyase-like, and the three MGL groups—draw on the single PF01053 candidate pool and resolve to three occupied nearest-reference categories, so the 23 target enzyme groups report as 20 reporting categories.

**Table 2.** Candidate counts for the 20 reporting categories, with the crosswalk from the 23 target enzyme groups in Table 1. The six PF01053 target enzyme groups resolve to three occupied reporting categories; no candidate was assigned nearest the cysteine-S-conjugate beta-lyase-like group, so that target enzyme group has no reporting category.

| Biochemical context | Target enzyme group | Reporting category | Pfam profile or architecture | Candidates |
| --- | --- | --- | --- | --- |
| Sulfur amino-acid and glutathione metabolism | OASTL | PF00291 pyridoxal phosphate-dependent enzyme domain | PF00291 | 33 |
| Sulfur amino-acid and glutathione metabolism | SAMS | S-adenosylmethionine synthase | PF00438 + PF02772 + PF02773 | 2 |
| Sulfur amino-acid and glutathione metabolism | GSH1 | Glutamate-cysteine ligase | PF04107 | 2 |
| Sulfur amino-acid and glutathione metabolism | CBL | PF01053 nearest-reference CBL | PF01053 | 2 |
| Sulfur amino-acid and glutathione metabolism | CGS | PF01053 nearest-reference CGS | PF01053 | 2 |
| Sulfur amino-acid and glutathione metabolism | GSH2 | Glutathione synthetase | PF03199 + PF03917 | 1 |
| Sulfur amino-acid and glutathione metabolism |  | **Context subtotal** |  | **42** |
| Conjugation, transfer, and redox reactions | UGT | UDP-glycosyltransferase domain | PF00201 | 218 |
| Conjugation, transfer, and redox reactions | GST | GST_N domain | PF02798 | 83 |
| Conjugation, transfer, and redox reactions | FMO | FMO-like domain | PF00743 | 40 |
| Conjugation, transfer, and redox reactions | SULT | SULT domain | PF00685 | 22 |
| Conjugation, transfer, and redox reactions | Rhodanese | Rhodanese domain | PF00581 | 16 |
| Conjugation, transfer, and redox reactions | GGT | GGT domain | PF01019 | 5 |
| Conjugation, transfer, and redox reactions |  | **Context subtotal** |  | **384** |
| Sulfur release and related reference pathways | Myrosinase | glycoside hydrolase family 1 (GH1) beta-glucosidase | PF00232 | 44 |
| Sulfur release and related reference pathways | Lachrymatory-factor synthase | Polyketide_cyc2 domain | PF10604 | 8 |
| Sulfur release and related reference pathways | Alliinase | Alliinase/ACC-synthase-like | PF04863 + PF04864 | 5 |
| Sulfur release and related reference pathways | MGL-Artocarpus, MGL-Cannabis, MGL-Durio | PF01053 nearest-reference MGL | PF01053 | 2 |
| Sulfur release and related reference pathways | CSCBL-like | no reporting category | PF01053 | 0 |
| Sulfur release and related reference pathways |  | **Context subtotal** |  | **59** |
| Oxylipin and volatile-product biosynthesis | AOS/HPL-motivated P450 search | P450 domain | PF00067 | 359 |
| Oxylipin and volatile-product biosynthesis | ADH | ADH architecture | PF00107 + PF08240 | 61 |
| Oxylipin and volatile-product biosynthesis | LOX | LOX architecture | PF00305 + PF01477 | 30 |
| Oxylipin and volatile-product biosynthesis | AOC | AOC domain | PF06351 | 6 |
| Oxylipin and volatile-product biosynthesis |  | **Context subtotal** |  | **456** |
|  |  | **Total** |  | **941** |

### 3.5 Expression mapping linked 168 candidates to 128 atlas genes

BLASTp returned 1,062 HSPs for 900 of the 941 candidates; 41 candidates had no alignment. At 100% identity across at least 80% of the GMO query, 168 of 941 candidates (17.9%) mapped to the Cannabis Expression Atlas, corresponding to 128 distinct atlas subject identifiers (Figure 4A). Mapping counts increased to 440 candidates at 99% identity, 584 at 98%, 695 at 95%, 795 at 90%, and 860 at 80%, all with query coverage of at least 80%.

**Figure 4.**
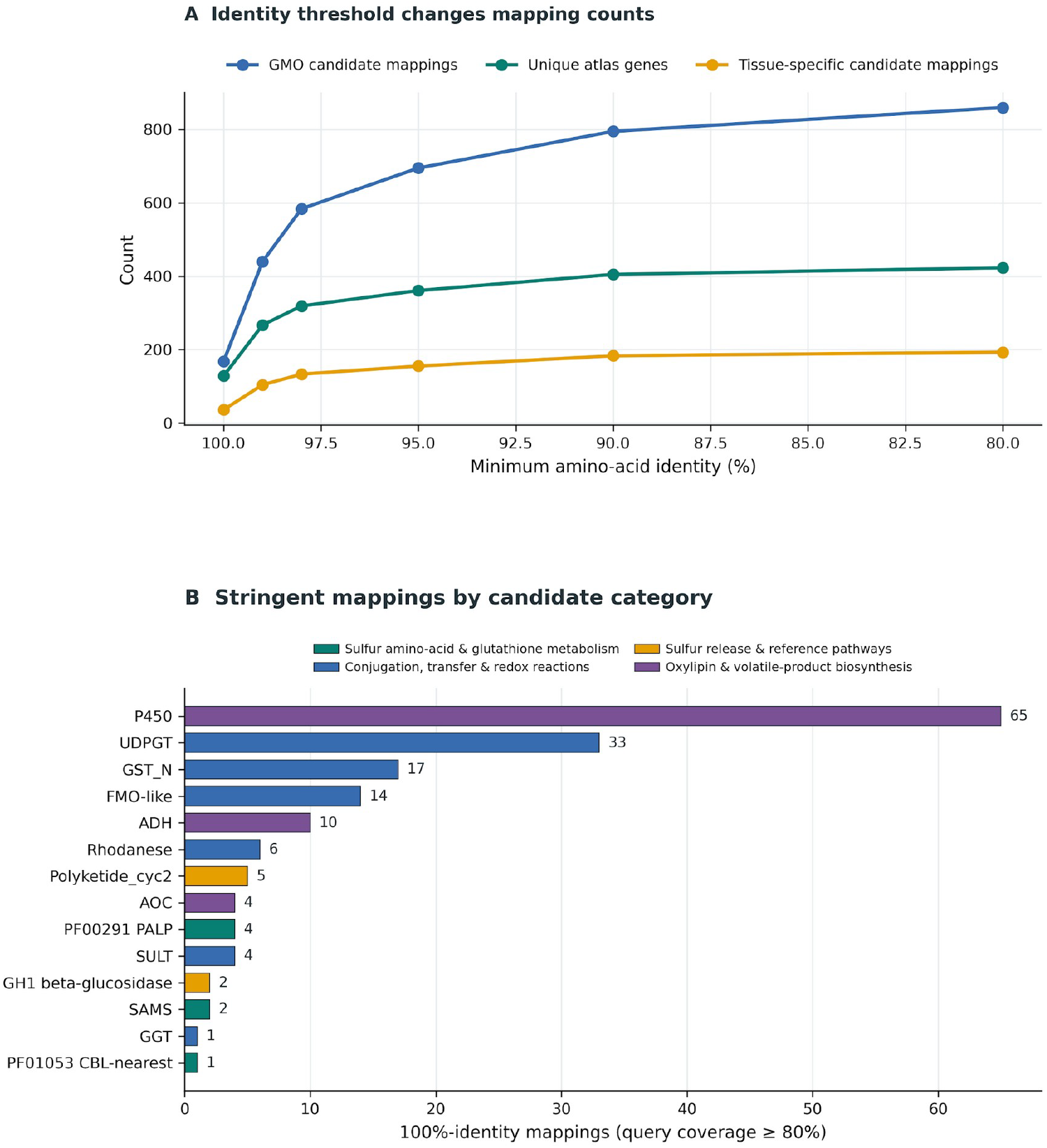
Identity-threshold sensitivity of Cannabis Expression Atlas mapping. **(A)** Candidate, distinct-gene, and tissue-specific mapping counts across 80–100% amino-acid identity at ≥80% query coverage. **(B)** Category composition of the 100%-identity set: 168 candidate mappings to 128 atlas genes, including 29 tissue-specific genes. In panel B, colors denote sulfur amino-acid and glutathione metabolism (teal), conjugation, transfer, and redox reactions (blue), sulfur release and related reference pathways (orange), and oxylipin and volatile-product biosynthesis (purple).

The mappings at 100% identity and at least 80% query coverage represented 14 of the 20 candidate categories (Figure 4B). The largest candidate-mapping counts were 65 for the P450-domain category, 33 for the UDP-glycosyltransferase-domain category, 17 for the GST_N-domain category, and 14 for the FMO-like-domain category.

Among the 128 distinct atlas genes, 41 were classified as expressed in all tissues, 33 as mixed, 29 as tissue-specific, 18 as low expressed, 6 as group enriched, and 1 as housekeeping. The tissue-specific genes comprised 13 root, 6 trichome, 5 hypocotyl, 2 seed, 1 induced male flower, 1 bast fibre, and 1 stem gene. At the candidate-mapping level, 36 mappings were tissue-specific, including 14 root-specific and 8 trichome-specific mappings.

## 4. Discussion

### 4.1 A broad candidate landscape for Cannabis sulfur-volatile research

Filtering retained 941 proteins across 20 sequence-defined candidate categories (Table 2; Figure 2B). Category sizes ranged from two GSH1 candidates and one GSH2 candidate to 359 P450-domain candidates, and the four largest categories accounted for 721 of the 941 proteins. This range provides both broad and narrow entry points for further investigation. The larger families expose potential sequence diversity but require substantial refinement, whereas the smaller groups provide more specific starting points for functional characterization. Because the counts also reflect the breadth of each search definition and the size of the underlying protein family, they do not necessarily show which categories contribute most to sulfur-volatile production.

The 359-member PF00067 set illustrates why broad families require additional resolution. The PF00067 domain alone cannot assign allene oxide synthase or hydroperoxide lyase activity. A Cannabis oxylipin inventory identified six CYP74 candidates and used focused phylogenetic analysis to support likely AOS or HPL activity for two of them (Borrego et al., 2023). The 13 proteins recovered by at least one CYP74 reference search should therefore be aligned at full length, placed in a focused phylogeny, and inspected for catalytic features before proteins are selected for assays.

Full-length-reference searches contributed 25 retained candidates that were not recovered by the corresponding Pfam profiles, including 24 UGT and one SULT candidate (Section 3.2). This suggests that more specific sequence queries can recover plausible candidates missed by domain-based searches and can help resolve large domain families.

The curated GMO v1 candidate set also provides a basis for extending these searches to other Cannabis proteomes. After refinement, the retained hits provide both individual high-confidence proteins for full-length similarity searches and alignments of well-supported homologs from which Cannabis-specific HMM profiles can be built. Full-length queries can identify close matches to specific candidates, whereas HMMs capture conserved sequence patterns across a homolog set and can identify more divergent family members. Applying both approaches to other cultivars and pangenome haplotypes could be used to distinguish candidates that are broadly conserved from those that vary in sequence or presence among cultivars and map the overall genetic diversity of these genes.

In general, these analyses provide a route from the broad catalog to progressively more specific targets. Following initial candidate selection and sequence screening, broad categories can be reduced further through full-length alignment, focused phylogeny, and inspection of catalytic features. Once candidate lists are narrowed, computationally intensive pipelines such as structure prediction and docking, which are impractical across the full search space, become feasible for further characterization and prioritization. Paired with sulfur-aroma phenotypes, this diversity map could support genotype–phenotype testing, marker development, and selective breeding.

### 4.2 Comparative systems define testable biochemical hypotheses

The candidate catalog supports several biochemical hypotheses, but homology and sequence screening alone cannot establish which candidates participate in sulfur-volatile production. The most informative first step is to measure candidate expression, relevant metabolites, and sulfur volatiles in the same samples. The comparative systems below identify the substrates and reactions to test.

One testable model for Cannabis C6 sulfur volatiles links sulfur amino-acid metabolism with glutathione synthesis and processing, as suggested by grapevine and wine-yeast biochemistry. Arabidopsis GSH1 and GSH2 have established roles in glutathione biosynthesis (Ullmann et al., 1996; Jez et al., 2004), while studies in grapevine demonstrate glutathionyl precursor formation and gamma-glutamyl processing for C6 thiols (Kobayashi et al., 2011; Philips et al., 2019). The corresponding GSH1, GSH2, GST, GGT, and pyridoxal-phosphate-dependent candidates should therefore be evaluated alongside targeted measurements of glutathionyl or cysteinyl precursors and the matching volatile in the same tissue. Substrate-specific enzyme assays should follow when these metabolites and candidate expression co-occur.

Additional candidate sets motivate tests of sulfur-cleavage reactions. These include five PF04863+PF04864 proteins with alliinase/ACC-synthase-like architecture and 44 PF00232 proteins with a GH1 beta-glucosidase domain (Table 2), as well as two MGL-nearest PF01053 proteins (Figure 3). These sequence features do not show that Cannabis uses alliinase-like or myrosinase-like pathways. Their relevance should first be evaluated by searching Cannabis tissues for compatible sulfur-containing substrates.

The PF01053 comparison also constrains this sulfur-cleavage hypothesis. No candidate resolved nearest the cysteine S-conjugate beta-lyase-like references, although that was the largest PF01053 reference group (17 of 41 entries) and contained four of the seven reviewed reference proteins. This result limits support for the specific Irc7-analogous hypothesis within the configured comparison, but it does not exclude beta-lyases outside the PF01053 profile.

For the C6 branch, the oxylipin candidates address a different question: formation of the upstream carbonyl skeleton. Hydroperoxide lyase can generate C6 aldehydes, whereas allene oxide synthase directs fatty-acid hydroperoxides toward jasmonate biosynthesis (Song et al., 1993; Wan et al., 2013; Borrego et al., 2023). The LOX, P450-domain, AOC, and ADH candidates provide entry points for testing whether Cannabis C6 sulfur volatiles share upstream carbonyl metabolism with other plant volatile pathways. Measurements across development and after biotic or abiotic challenge could distinguish candidates associated with sulfur aroma from proteins with broader stress or oxylipin roles.

The sequence basis of the prenyl carbon-sulfur linkage remains unresolved. Experiments that supply Cannabis tissues with candidate precursor compounds and use isotope labels to track those compounds through metabolism could help identify the source of the prenyl carbon skeleton and determine when sulfur is introduced or released.

### 4.3 Expression mapping narrows candidates for follow-up

Among the 29 tissue-specific genes in the 100%-identity mapping set, 13 were root-specific and 6 were trichome-specific (Figure 4). Trichome-specific expression provides a reason to prioritize these genes for aroma-focused testing, whereas root-specific genes may support general sulfur or root metabolism rather than floral aroma. Matched expression and sulfur-volatile measurements in roots, trichomes, and flowers are needed to distinguish these roles.

Mapping counts increased from 168 candidates at 100% identity to 860 at 80% identity (Figure 4A). However, this remains a screening map rather than functional validation. Several pipeline limitations support this interpretation: the analysis covers one GMO v1 proteome; broad domain and nearest-reference matches do not establish catalytic activity; tree assignments were not filtered by branch support; and motif screening did not narrow the catalog. Rather than providing direct functional validation, the 100%-identity mappings across at least 80% of each query define the first experimental-priority tier. Lower-identity matches provide broader expression evidence for homologs rather than equivalent gene-level links, and their counts are lower bounds because BLASTp retained one target per query. More specific full-length protein queries are needed to refine candidate identification. Additional ribonucleic acid sequencing (RNA-seq) paired with sulfur-volatile measurements in roots, trichomes, and flowers is needed to refine tissue prioritization.

## 5. Conclusion

This study narrowed the search for Cannabis sulfur-volatile enzymes from 55,790 proteins to 941 candidates across 20 sequence-defined reporting categories. By integrating profile and full-length reference searches, sequence screening, focused phylogenetic assignment, and expression mapping, the analysis converted an open-ended proteome-scale question into a structured set of sequence-defined candidate groups and biochemical hypotheses. Mappings at 100% HSP identity and at least 80% query coverage linked 168 candidates to 128 Cannabis Expression Atlas genes, including 29 with tissue-specific expression, further refining targets for future experimentation.

The catalog separates C6 and prenylated sulfur volatiles into distinct biochemical problems. For the C6 branch, candidates related to glutathione processing, sulfur release, and formation of the upstream carbonyl skeleton can be tested through matched measurements of expression, precursors, volatiles, and enzyme activity. The sequence basis of the prenyl carbon-sulfur linkage remains unknown; experiments that supply Cannabis tissues with candidate precursor compounds and use isotope labels to track those compounds through metabolism could help identify the source of the prenyl carbon skeleton and determine when sulfur is introduced or released. Across the catalog, trichome-specific genes are candidates for floral-aroma studies, whereas root-specific genes require broader interpretation.

The resulting panel provides candidate proteins for follow-up studies of Cannabis sulfur-aroma biosynthesis. Building on this foundation, searches with additional reference proteins and genomes can refine the panel and reveal variation across cultivars and pangenome haplotypes. Additionally, protein-structure modeling and docking can generate testable predictions of enzyme function and substrates. These predictions can then be validated with RNA and protein measurements, enzyme assays, and measurements of pathway intermediates and volatiles. Together, this work can define sulfur-aroma pathways, identify useful genetic variation, and guide selective breeding.

## Data availability

The supporting dataset, including query definitions, search assignments, staged validation results, retained candidate sequences, expression cross-references, and PF01053 phylogenetic artifacts, is archived on Zenodo (version 0.1.0; https://doi.org/10.5281/zenodo.21685901). The GMO assembly and its version 1 gene annotation are available through the pangenome resources reported by Lynch et al. (2025), and Cannabis Expression Atlas resources are available through Barbosa-Xavier et al. (2024).

## AI-assisted manuscript preparation

Large language models were also used to assist with manuscript drafting, restructuring, consistency checking, and proofreading. All model outputs were reviewed and edited by the authors, who made all final scientific and editorial decisions and take responsibility for the manuscript’s accuracy and integrity.

